# Northern Cerrado Native Vegetation is a Refuge for Birds Under Current Climate Change

**DOI:** 10.1101/2021.11.17.468959

**Authors:** José Hidasi-Neto, Nicole Mércia Alves Gomes, Nelson Silva Pinto

**Affiliations:** No affiliation, Goiânia, Brazil; Programa de Pós-Graduação em Biodiversidade Animal, Universidade Federal de Goiás, Goiânia, Brazil

**Keywords:** Avifauna, ecological niche modelling, conservation, biotic homogenization

## Abstract

Climate Change is already seen as one of the biggest threats to biodiversity in the 21^st^ century. Not much studies direct attention to its effects on whole communities of threatened hotspots. In the present work, we combine ecological niche modelling (ENM) with a future climate scenario of greenhouse gases emissions to study the future changes in alpha and beta diversity of birds of the Brazilian Cerrado biome, a hotspot of biodiversity with high velocity of climate change and agricultural expansion. In general, we found heterogeneous results for changes in species richness, spatial and temporal taxonomic and functional beta diversity, and mean ecological distinctiveness. Contrary to a previous study on Cerrado mammals, species richness is expected to increase in Northern Cerrado, where homogenization of communities (decreasing spatial turnover) is also expected to occur especially through local invasions. We show that biotic homogenization (which is composed of local extinction of natives and local invasion of exotic species) will occur in two biological groups but through different subprocesses: local extinctions for mammals and local invasions for birds. Distinct conservation management actions should be directed depending on the outcomes of analyzes of alpha and spatial and temporal beta diversity, for example controlling species invasions in Northern Cerrado. Conservation studies should continue evaluating Cerrado in Brazil even under covid pandemic, as environmental situation in the country is not good and incentives for scientific studies are almost nonexistent.

## Introduction

Earth is facing a, most probably anthropic, climate change, which is threatening great part of biodiversity in the 21^st^ century [1]. Although United Nations’ Intergovernmental Panel on Climate Change (IPCC) has declared several climate change scenarios in relation to the emission of greenhouse gases (from worst to optimistic), we can expect changes to be bad for species, but not always expecting the worst scenario, for this scenario “becomes increasingly implausible with every passing year” [2,3]. Even so, the synergistic effects of climate change and land-use is leading biological groups to severe impacts [4]. It is also notable that climate change does not threaten biomes equally. The velocity of climate change can be different depending on regional climatic features. Mountainous biomes show the slowest velocity of climate change, while flatter biomes such as mangroves, flooded grasslands and savannas show the highest velocities [5]. Great focus should be pointed to the latter ecosystems as they are the ones closest to a biodiversity collapse [6].

The Cerrado biome is not only a flatter biome, therefore presenting a high velocity of climate change [5], but also a hotspot of biodiversity [7]. In other words, it is both threatened and presents a high number of endemic species. Politically speaking, Cerrado is now facing a decisive moment to deal with drivers of extinction of its rich biodiversity, with conservation planners focusing on the conservation of areas with high amount of native vegetation (mostly Northern Cerrado) [8] while restoring sites which suffered habitat loss or fragmentation (mostly Southern Cerrado), but that can still harbor an expressive number of endemic species [9]. Summing up all this information about Cerrado, the biome goes to the priority list of regions to be studied in the current century. Indeed, the number of publications related to Cerrado biodiversity and conservation has immensely increased since the 90s [8], and it seems to continue increasing in the following years or even decades.

Studies on the effects of climate change on biodiversity usually focus on populations and species. Some species can benefit from climate change, increasing their abundances or widening their distributions, while others remain the same or suffer negative consequences [10,11]. However, climate change will most likely change the spatial distribution of environmental conditions faster than species are able to adapt [10,12,13], especially in a warm flat biome such as the Brazilian Cerrado. In consequence, we expect species richness of biological groups to decrease throughout the biome. Notably, some studies have showed the effects of climate change on whole communities [12–16]. It is expected a loss of species richness for biological groups, such as tropical plants and mammals [16,17]. This provides evidence that species from ecosystems such as tropical ones will not easily track changes in the spatial distribution of their environmental conditions.

Some studies on climate change effects on whole communities not only find evidence that alpha diversity will be changed, but also beta diversity [16,18], which can be mostly reduced. This is a great conservation concern as these findings can be related to a process of homogenization occurring on fauna and flora throughout the world [19–21]. A study with bird communities in Mexico presents the possibility of biotic homogenization resulting from anthropogenic interventions in landscape [22]. Human interventions also can lead to taxonomic and functional homogenization in bird communities [23], in addition to influencing the reduction of local richness and species abundances. Currently, the north portion of Cerrado, although it is widely used for farming purposes, is still more vegetated than the south portion. The synergetic effects of human interventions and climate change can become potential threats for local biodiversity, in view of potential endemism in this biome. Biotic homogenization is a process which combines two subprocesses: the extinction of native specialist species, and the invasion of exotic generalist species [21,24,25]. In a study on Great Britain, biotic homogenization was a predicted factor as one of the effects of anthropogenic environmental changes in bird communities, indicating that rising temperature can lead to decreased specialization [26]. There are other factors that determine homogenization in addition to species introductions and extinctions, such as the evolutionary history of groups and the spatial scale [27]. It has been found that climate change can homogenize mammalian communities in Southern Cerrado in the near future, mainly due to the local extinction of species in a region with little native vegetation [16]. Similarly, we would expect communities from different biological groups to become more similar among each other in the future, especially in regions with low amount of native vegetation. Moreover, as a result of the subprocesses of biotic homogenization, we expect homogenized regions to present communities with low mean of species ecological originality, as ecologically original species become locally extinct while ecologically non-original species arrive from outside regions.

In the present work we project the potential distributions of the birds of Cerrado, while analyzing the alpha, spatial beta, temporal beta, and mean ecological distinctiveness of species. We also discuss our findings in light of previous research regarding the Cerrado biome. Specifically, we answer the following questions: (1) Are we losing Cerrado bird species? (2) Are bird communities becoming more homogenized with time? (3) How and where are these changes going to happen in the biome? (4) Are ecologically original species being replaced by ecologically non-original species in areas that will become more biotically homogenized?

## Materials and Methods

### Study Area

We studied birds from the Cerrado biome, which covers approximately 2,000,000km^2^. The biome has a very heterogeneous vegetation, presenting grassland savannas, semideciduous forests, wet grasslands and rupestrian fields. In general, it presents a humid tropical climate, showing wet summers and dry winters [28]. It is considered to be one of the most threatened regions of the world, due mainly to agriculture-related habitat destruction and fragmentation [28].

### Climate, Species Occurrence, and Trait Data

First, we downloaded the present (or historical) and future climate bioclimatic variables from WorldClim 2.1 (2.5 minutes) [29]. The future data was for the 2081-2100 period under SSP (Shared Socioeconomic Pathway) 245 (intermediate concentration of greenhouse gases), for both MIROC6 (Model for Interdisciplinary Research on Climate) and MRI (Meteorological Research Institute) General Circulation Models (GCMs) [30]. We intersected these climatic scenarios with a shapefile of South America. We then got a list of birds from South America. To do so, we downloaded the distributions of all birds of the world [31] and intersected them with a shapefile of South America. Then, we used this list to get the occurrence points for each species from Global Biodiversity Information Facility (GBIF) [32]. We removed species with less than 25 occurrences. Following, we created a grid (0.5° × 0.5°) from a Cerrado shapefile, removing cells with less than 50% cell area. We resampled means of bioclimatic variables from our three (present and two GCMs) climate scenarios using this Cerrado grid. We got trait data from EltonTraits 1.0 [33] while fixing synonyms (but see how species will be considered or removed from analyses in the following subsection).

### Ecological Niche Modeling

For each species of our South American bird list, we used Ecological Niche Modeling (ENM) to observe if it occurs in the present-day or future Cerrado. To do that we used occurrences and climate data from South America to calculate GLM (General Linear Model) and Maxent (Maximum Entropy) algorithms, using 20% - 80% test-training procedure, and using 1000 random pseudo-absences as background. As different results could come from the GCMs and algorithms, it is recommended to use ensemble forecasting to merge results based on TSS (True Skill Statistics) [34,35]. Then, for the present we did an ensemble using GLM and Maxent results using the Cerrado grid with climatic data as a focus of study, while for the future we calculated ensembles for these algorithms for both MIROC and MRI GCMs, ending up with mean habitat suitability values from both future ensembles (again using Cerrado as a focus of study). To perform ENMs and ensembles we used “sdm” package in R software [36,37]. We used a congruence threshold higher than 0.5 to obtain a final presence/absence map for each species (habitat suitability > 0.5 as presence). In that manner we came up with bird occurrence matrices for both present and future Cerrado.

### Analyses

In order to observe if climate change will reduce species richness, we first used the occurrence maps to calculate richness in both the present and the future, calculating the percentage change in relation to present-day Cerrado. Then, to observe if climate change will generate biotic homogenization in Cerrado, for each focal cell we calculated mean spatial turnover in relation to its 8 neighbor cells (some cells presenting less than 8 neighbors). We did that in both the present and future, calculating also the change of future-present in values of spatial turnover [16]. We also calculated the temporal turnover and nestedness between the present and future Cerrado to be able to analyze regional changes due to richness or composition change. We used occurrence matrices and trait data to filter Cerrado’s present-day and future species and create a functional dendrogram. To do that we used data on body mass (quantitative), activity period (binary), diet (fuzzy) and forage stratum (fuzzy), calculating a Gower’s distance matrix, then a UPGMA (Unweighted Pair Group Method with Arithmetic Mean) functional dendrogram. We used this dendrogram to calculate spatial functional turnover (same as spatial taxonomic turnover noted above, but considering a functional dendrogram), using the turnover partition of PhyloSor index [16,38]. We also used the same index to calculate temporal functional turnover and nestedness. Finally, to analyze if ecologically non-distinct birds arrive at places with ecologically distinct birds, reducing local or regional mean ecological distinctiveness, we also used the functional dendrogram to calculate ecological distinctiveness (EcoD; evolutionary distinctiveness but using a functional dendrogram), using the fair proportions method [39,40]. We calculated mean EcoD for each grid cell in both the present and the future and calculated the percentage change in mean EcoD in relation to the present.

## Results

We observed that, in general, there will be a reduction of bird species richness (−6.40 ± 23.465 species). We will have 237 bird regional extinctions and 116 regional invasions (see Supplementary Material for more details). We observed a heterogeneous pattern of climate change effects on local species richness (Figure 1). While we observed a reduction in richness throughout Southern Cerrado, there was an increase in richness in Northern Cerrado in the future.

**Figure 1.**
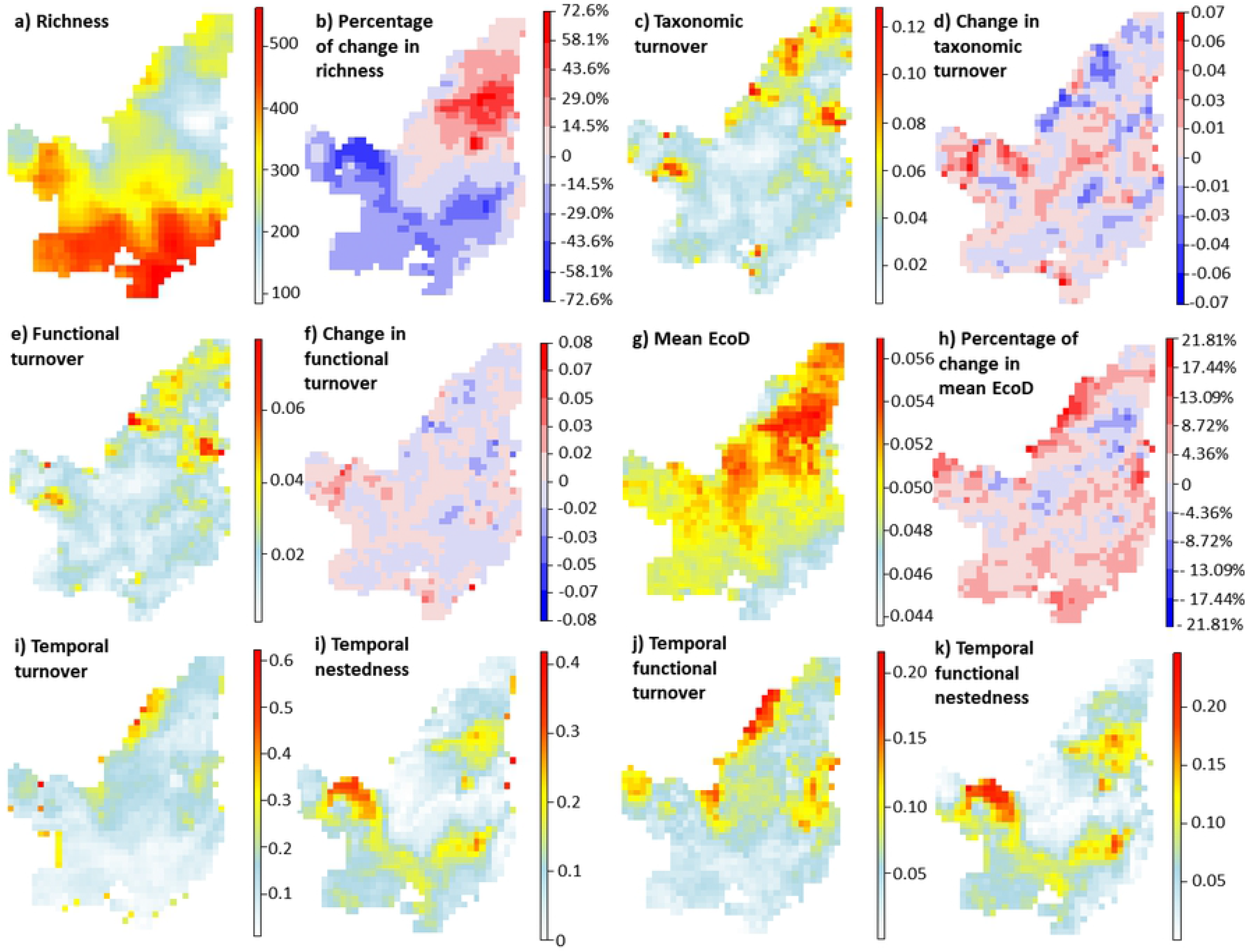
Maps indicating the present values and the future change in richness (“a” and “b”), spatial taxonomic turnover (“c” and “d”), spatial functional turnover (“e” and “f”), mean EcoD values (“g” and “h”) between present time and the 2081-2100 period under a scenario of intermediate concentration of greenhouse gases (SSP 245). There are also temporal turnover and nestedness (“i” and “j”) and temporal functional turnover and nestedness (“j” and “k”) between the same periods of time.

We then observed spatial and temporal turnover and nestedness. Even if heterogeneous, we observed a decrease in spatial taxonomic and functional turnover in Northern Cerrado. Curiously, this pattern was linked with low temporal taxonomic and functional turnover and high temporal taxonomic and functional nestedness. So, this observable pattern is related to an influx of species in Northern Cerrado, homogenizing communities mainly due to local invasions.

Finally, looking at mean Ecological Distinctiveness (EcoD) values, we observed that an increase in species richness, decrease in spatial taxonomic and functional turnover, low temporal taxonomic and functional turnover, and high temporal taxonomic and functional nestedness was related to a decrease in mean EcoD. So, it is observable an influx of non-original species in future Northern Cerrado, decreasing the mean value of distinctiveness of communities.

## Discussion

We observed a heterogeneous pattern of results for changes in alpha and beta diversity of birds in the Brazilian Cerrado. We found highest bird species richness in Southern Cerrado, while we expect Southern Cerrado to greatly lose, and Northern Cerrado to greatly increase species richness. Northern Cerrado is expected to show considerable reduction in spatial taxonomic and functional turnover, while high temporal taxonomic and functional turnover and mostly low temporal taxonomic and functional nestedness. Communities are also expected to become less ecologically original (low mean EcoD). Our results were, in part, contrary to what was found for mammals by Hidasi-Neto et al. [16], where homogenization was, in turn, higher in Southern Cerrado. Summing up all results from this present work and Hidasi-Neto et al., Northern Cerrado, mainly due to its high amount of native vegetation remaining (Figure 2), is a refuge for Cerrado mammal and bird species, but presenting different subprocesses regulating the homogenization of communities.

**Figure 2.**
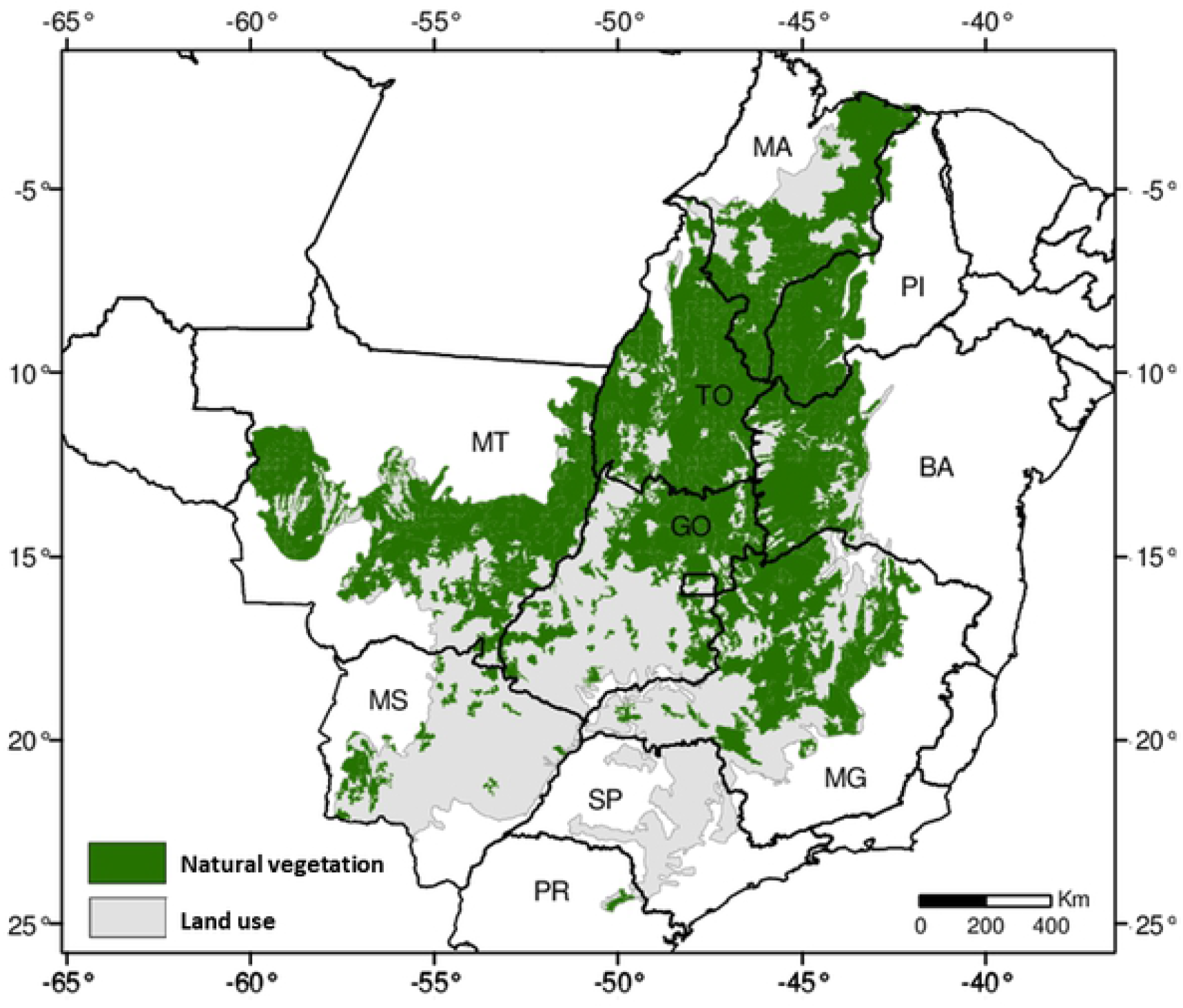
Cerrado natural vegetation and land use regions. Figure modified from [41,42].

Bird species richness is highest in Southern Cerrado, where there will also be a loss of species richness in the future. On the other hand, Northern Cerrado is expected to receive species, due to local invasions. This result is somehow different to those found for mammals of Cerrado [16], which are expected to reduce throughout all the biome. Moreover, protected areas (PAs) represent only 8.3% of Cerrado, or 6.5% if only considered the fraction covered by native vegetation [8]. So, PAs are not able to conserve species richness of biological groups, and, in the case of mammals, they are not able to conserve mammal evolutionary history and trait diversity [43]. Notably, in a recent study on the refugia for Cerrado birds, 13% of Cerrado was observed with high native vegetation and low climatic anomaly (magnitude of change in the mean climatic conditions) [44]. These areas are mainly in the Eastern to Northeastern part of Cerrado [44], and although only 15% of studied bird distributions fall within this fraction of Cerrado (refugia are not species-rich areas), these are stable regions to conserve birds in the biome. Based on our results, and in accordance with Hidasi-Neto et al. [16], more attention should be focused on Northern Cerrado, which today presents the highest amount of native vegetation in the biome [41,42], and also where agricultural expansion is likely to take place in the coming years [45].

Based on our results and on Hidasi-Neto et al. [16], we observed different patterns of biotic homogenization throughout Cerrado. In Hidasi-Neto et al., Southern Cerrado was mainly homogenized in relation to it mammal communities. Contrary to this finding, we observed in this present work a biotic homogenization mainly in Northern Cerrado. However, it is notable that different processes cause these distinct patterns. While for mammals, local extinctions were mainly responsible for the homogenization in Southern Cerrado (Figure 3), for birds, local invasions were responsible for the homogenization in Northern Cerrado (Figure 3). Indeed, local invasions (alongside deforestation and wildfires) are expected to occur in areas that today present high amount of native vegetation [44]. Considering our results of alpha and taxonomic and functional beta diversity, it is urgently needed a integrative approach to identify and conserve species and communities focusing not only on taxonomic richness and composition, but also on trait diversity related to ecosystem functions and services, while regenerating Southern Cerrado and promoting ecosystem connectivity [40,43,46].

**Figure 3.**
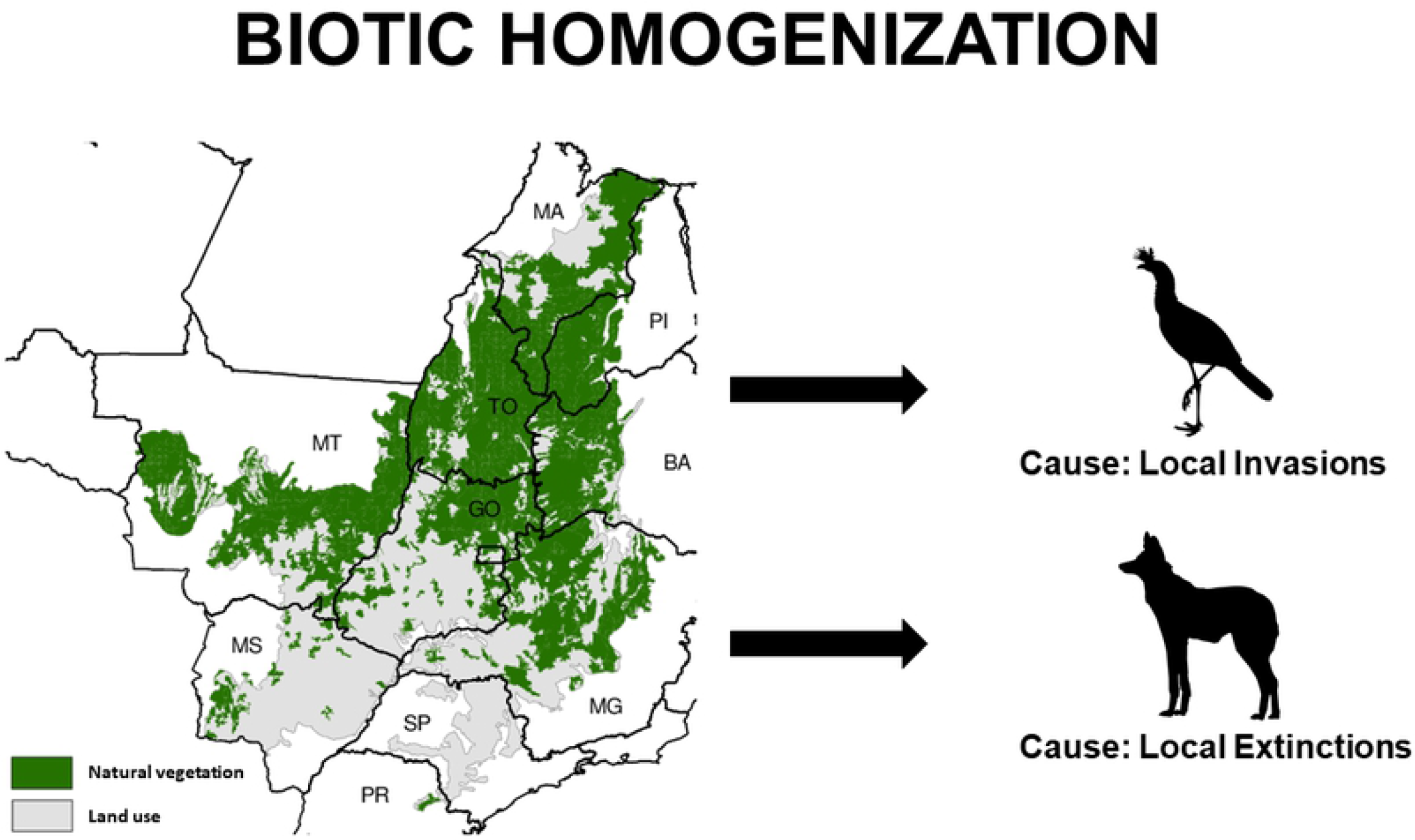
Biotic homogenization in Cerrado biological groups and its causes.

Results are not surprising if you already know how native vegetation covers present-day Cerrado (Figure 2). However, it is surprising that different subprocesses (local extinction vs local invasions) can generate distinct projections on how climate change will modify communities. As already stated, native vegetation is mainly found in Northern Cerrado. Cerrado native vegetation is mainly threatened by agricultural expansion [8,44,45]. As showed in Figure 1 in Sobral et al. [47], there are different possible findings when you analyze the alpha and temporal beta diversity of communities. This relationship is also addressed in Hidasi-Neto et al. [16]. For example, we expect to see a ‘compensation and gain’ in Northeastern Cerrado, where part of refugia are expected to be found [44]. In accordance with past studies [8,44], we propose different conservation management actions to be used depending on the expected outcome in a focal Cerrado area considered for conservation. Present and past results for birds and mammals come to a consensus that Northern Cerrado’s native vegetation should be conserved for being a future refuge for Cerrado species.

The Cerrado biome presents a heterogeneous vegetation, with several phytophysiognomies that cover both forest and non-forest natural areas. However, most conservation efforts focus on forest remnants, disregarding the heterogeneity of Cerrado vegetation at expense of natural open fields, which are more targeted for farming purposes [48]. The private lands concentrate the most part of native forest remnants of Brazil [49], generally in Legal Reserves (LR), protected areas for the law, that are selected by the owners and are often the least viable areas for agricultural use. In the Cerrado biome, the LR should correspond to 35% of land that is part of the Legal Amazon, and only 20% in the rest of biome [50], and the non-forest areas are the less protected inside private properties [48]. The open areas of the Cerrado have already been identified as important for maintaining the functional diversity of bird communities [51] and for diversity of anurans [52]. Thus, future conservation and restoration plans must be designed in such a way as to maintain the plant heterogeneity of the Cerrado and consider the diversity indices of endemic species to conserve biodiversity and reduce impacts on Cerrado bird communities.

Computational resources are being increasingly available to common people. Methods that could take several months in workstations are now feasible within weeks. This means a lot to studies on climate change effects on communities, especially if resources are limited due to special cases, such as covid pandemic. This shows that conservation planning is able to develop itself and continue in difficult times. Brazil, as other countries, is having a difficult time in relation to its environmental conservation, and Cerrado is a highly endangered hotspot of biodiversity that faces agricultural-based habitat loss and fragmentation. Efforts should be directed in the following years to protect Cerrado’s native vegetation, while recovering vegetation in Southern Cerrado.

## Acknowledgments

We thank the anonymous reviewers for critical comments on the text. We also thank SUS (Sistema Único de Saúde) for the battle against covid in these mostly dark times.

## SUPPLEMENTARY MATERIAL CAPTION

(S1) Species occurring in the present and in the future, and whether they are a regional extinction or a regional invasion.

